# Explainable Active Learning Framework for Ligand Binding Affinity Prediction

**DOI:** 10.64898/2025.12.17.694851

**Authors:** Satya Pratik Srivastava, Rohan Gorantla, Sharath Krishna Chundru, Claire J.R. Winkelman, Antonia S.J.S. Mey, Rajeev Kumar Singh

## Abstract

Active learning (AL) prioritises which compounds to measure next for protein–ligand affinity when assay or simulation budgets are limited. We present an explainable AL framework built on Gaussian process regression and assess how molecular representations, covariance kernels, and acquisition policies affect enrichment across four drug-relevant targets. Using recall of top active compounds, we find that dataset identity—the target’s chemical landscape—sets the performance ceiling, while method choices modulate outcomes rather than overturn them. Fingerprints with simple Gaussian process kernels provide robust, low-variance enrichment, whereas learned embeddings with non-linear kernels can reach higher peaks but with greater variability. Uncertainty-guided acquisition consistently outperforms random selection, yet no single policy is universally optimal; the best choice follows structure-activity relationship (SAR) complexity. To enhance interpretability beyond black-box selection, we integrate SHapley Additive exPlanations (SHAP) to link high-impact fingerprint bits to chemically meaningful fragments across AL cycles, illustrating how the model’s attention progressively concentrates on SAR-relevant motifs.

## 1 Introduction

Drug discovery is the process of identifying new molecules that can target a disease state with novel chemical compounds ranging from small molecules [1–3] to anti-bodies [3, 4]. One way of approaching small molecule drug discovery is by identifying a biological target, e.g., a protein or other relevant biomolecule to alter their functional state by inhibition. One key property that can help identify novel inhibitors for a protein target is optimisation of the protein-ligand binding affinity. As such, accurate *in-silico* and experimental estimation of protein–ligand binding affinities are essential properties to measure and predict during hit identification across vast chemical libraries and systematic optimization of congeneric series during hit–to–lead campaigns [5–7]. High–throughput screening remains a cornerstone of small-molecule discovery, but rising assay complexity and cost increasingly preclude exhaustive use [8]. As discovery shifts toward medium–throughput, biophysics-rich assays supported by structure-guided optimisation with alchemical free-energy methods (AFE) [2, 6, 9–11], the goal centers around exploring chemical space effectively under a budget. This budget can determine both the number of evaluations measurements or computational predictions by only assessing a few hundred compounds per cycle [5, 6, 12]. Biophysical assays such as surface plasmon resonance (SPR) provide kinetics-resolved confirmation of binding and are widely used when functional assays are noisy, non-specific, or fail to identify tractable series [13, 14]. Yet their medium-to-low throughput constrains campaign scale. Similarly, AFE calculations can prospectively prioritise substitutions but remain computationally intensive. In both cases, identifying the most promising set of compounds with the fewest computational or experimental evaluations and minimising the overall budget is desirable.

Active learning (AL), a subset of machine learning, has emerged as a framework to address this challenge [2, 9, 15]. By training a surrogate model, quantifying predictive uncertainty, and iteratively prioritising the next most informative compounds, AL turns limited assays or compute into maximal information gain, improving enrichment while reducing reliance on brute-force screening [9, 15–18]. In practice, AL balances *exploitation* that is refining known high-activity scaffolds, against *exploration* that probes novel chemotypes that may unlock new structure-activity relationship (SAR). This trade-off is controlled by the acquisition strategy [19, 20]. As a result, AL has been deployed for ligand binding affinity prediction and multi-property lead optimisation under assay- or simulation-constrained budgets [2, 9, 15, 18, 20]. Notwithstanding its potential, AL is not a “onesize-fits-all” solution [15, 21]. Its performance is significantly dependent on a complex interplay of methodological choices, including the underlying machine learning model, the molecular representation, the kernel function, and the acquisition protocol [19, 21]. Outcomes vary with the chemical landscape of the library, the molecular representation, the surrogate model choice, and the acquisition protocol [15, 19, 21]. Moreover, surrogate models and representations from deep learning models can behave as “black boxes,” limiting chemical intuition and trust in recommendations [22–24]. AL has been applied successfully on individual targets [2, 25, 26] and specific workflows [10, 27, 28], and recent efforts have begun to systematically explore different strategies and parameters [15, 21]. Open questions remain around clarifying when different AL designs are most effective, why performance varies across chemical spaces, and finding ways to incorporate explainability into the selection process of the AL cycles to help with guiding design choices that can be experimentally verified.

In this work we combine explainability while exploring seven acquisition protocols with five Gaussian-process kernels and three molecular representations (ECFP4, MACCS, ChemBERTa) in a fixed budget-setting for pharmaceutically relevant targets taken from literature (TYK2, USP7, D2R, MPro). We show that dataset identity i.e., the target’s chemical landscape dominates achievable enrichment, and that representation–kernel choices trade off robustness (fingerprints with simple kernels) versus peak performance (learned embeddings with non-linear kernels) is an important decision factor. To move beyond black-box selection, we integrate SHapley Additive exPlanations [29] (SHAP) to map high-impact fingerprint bits to chemically interpretable fragments over AL cycles, revealing how model focus sharpens onto SAR-relevant motifs.

## 2 Methods

### 2.1 Active learning setup

Central to our active learning framework is a methodology that employs principles of Bayesian Optimization (BO) [30]. BO is an iterative strategy for optimizing black-box functions that are expensive to evaluate. It operates by building a probabilistic surrogate model of the objective function, which is then used to intelligently select the most promising points to evaluate next. In our framework, the surrogate model approximates the relationship between molecular structure and binding affinity across the chemical space of ligands. An acquisition function (AF) uses the model’s estimates and uncertainties to select the next batch of compounds for evaluation. The model is then updated with the new data, and the process is repeated. The ultimate goal of this iterative process is to find the compound with the highest affinity, as summarised by the objective in Eq. 1.

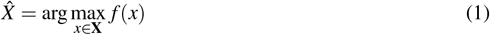

Here, *f* (*x*) represents the true but unknown binding affinity of a given compound (molecule) *x*. The search space *X* represents the entire library of candidate compounds available for evaluation. The goal of BO is to find the optimal compound 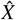 that maximizes the affinity, while minimizing the number of expensive evaluations of *f* (*x*) (i.e., experiments or simulations).

### 2.2 Gaussian Process as surrogate model

The most common and effective class of surrogate models for Bayesian Optimization are **Gaussian Processes (GPs)** [31]. A GP is a non-parametric model, defined by its mean function *m*(**x**) and covariance, i.e., kernel function *k*(**x, x**^*′*^) which measures the similarity between two points. The GP is defined by the following equation 2

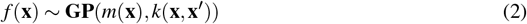

Gaussian functions can model the unknown affinity function **f**(**x**) on a distribution of functions, and they are incredibly adaptable at approximating nonlinear functions, which are needed to traverse the vast chemical space.

### 2.3 Acquisition strategies in AL cycles

Compound selection within the active learning loop is guided by an acquisition strategy; here we use the generalized Upper Confidence Bound (UCB) acquisition function [20, 32]. This function balances exploring new molecules with exploiting known good binders by linearly weighting the model’s estimated mean affinity and its associated uncertainty. The acquisition score for a compound *x* is determined as follows,

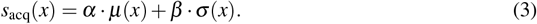

Here, *µ*(*x*) is the estimated mean affinity for *x, σ* (*x*) is the estimated standard deviation (uncertainty), *α* is a parameter weighting exploitation (mean prediction), and *β* is a parameter weighting exploration (uncertainty) [17]. Seven distinct acquisition strategies have been examined by varying *α* and *β* parameters over the acquisition cycles.

The seven distinct active learning acquisition protocols in this study were designed to systematically probe the trade-off between exploration and exploitation. Each protocol began with an initial random batch of 60 compounds to seed the model, followed by 10 acquisition cycles of 30 compounds each. The exploration-exploitation balance was controlled by dynamically varying the *α* and *β* parameters in the generalized Upper Confidence Bound (UCB) acquisition function:*s*_acq_(*x*) = *α* · *µ*(*x*) + *β* · *σ* (*x*).

This framework allows for three primary modes: **pure exploration** (*α* = 0, *β* = 1), which prioritizes molecules with the highest uncertainty (*σ* (*x*)); **pure exploitation** (*α* = 1, *β* = 0), which selects the most promising estimated affinity (*µ*(*x*)); and a **balanced strategy** (*α* = 0.5, *β* = 0.5). The specific schedules for each protocol are summarised in Table 1.

**Table 1.** Overview of Active Learning Acquisition Protocols. Each protocol starts with an initial batch of 60 randomly selected compounds, followed by 10 cycles of 30 compounds. **Shorthand:** R=Random, E=Explore (*α* = 0, *β* = 1), X=Exploit (*α* = 1, *β* = 0), B=Balanced (*α* = 0.5, *β* = 0.5). Numbers in parentheses indicate the number of compounds acquired in that step.

| Protocol Name | Acquisition Schedule (10 Cycles of 30 Compounds) |
| --- | --- |
| Random Baseline | $[R(30)] \times 10$ |
| UCB-Balanced | $[B(30)] \times 10$ |
| UCB-Alternate | $[E(30), X(30)] \times 5$ |
| UCB-Sandwich | $[E(30)] \times 2 + [X(30)] \times 6 + [E(30)] \times 2$ |
| UCB-Explore-heavy | $[E(30)] \times 7 + [X(30)] \times 3$ |
| UCB-Exploit-heavy | $[X(30)] \times 7 + [E(30)] \times 3$ |
| UCB-Gradual | $[E(30)] \times 3 + [B(30)] \times 4 + [X(30)] \times 3$ |

Beyond simple baselines like the Random and UCB-Balanced protocols, we designed several dynamic strategies to model different discovery campaign philosophies:

- **UCB-Alternate:** This protocol alternates every cycle between pure exploration and pure exploitation to explicitly separate the search for novel chemotypes from the refinement of known active scaffolds.
- **UCB-Sandwich:** This strategy “sandwiches” a long phase of intensive exploitation (6 cycles) between two short phases of initial and terminal exploration (2 cycles each), modeling a campaign that quickly focuses on a promising region before a final check for missed opportunities.
- **UCB-Gradual:** This protocol mimics a phased discovery campaign, beginning with broad exploration (3 cycles), transitioning to a balanced search (4 cycles), and concluding with focused exploitation (3 cycles) as the SAR landscape becomes better defined.

### 2.4 Model Validation and Hyperparameter Handling

To ensure the robustness of our models and the validity of their uncertainty estimates, we incorporated several validation and regularization techniques.

#### Hyperparameter Optimization and Regularization

Kernel hyperparameters and the model’s likelihood were optimized in each AL cycle by maximizing the marginal log-likelihood using the **Adam optimizer** for 100 epochs. To correct for potential model miscalibration, we implemented weakly informative **Gamma priors** on the GP model’s likelihood noise (Γ(1.1, 0.05)) and the kernel’s lengthscale parameter (Γ(3.0, 6.0)), a step proven to be critical for producing reliable uncertainty estimates (Supplementary Figure S.1).

#### Uncertainty Calibration Diagnostics

A core premise of UCB-based active learning is that the model’s predictive uncertainty, *σ* (*x*), is well calibrated. To validate this, we performed a suite of calibration diagnostics at the final cycle of each experiment. We calculated and analyzed three key metrics: **Probability Integral Transform (PIT) Histograms** to assess distributional correctness, **Reliability Diagrams** to check the accuracy of confidence intervals, and the **Negative Log Predictive Density (NLPD)** to provide an overall score for the predictive distribution.

#### Preprocessing Ablation Study for ChemBERTa

To investigate the sensitivity of non-Tanimoto kernels to the scale of high-dimensional ChemBERTa embeddings, we conducted a comprehensive **ablation study**. We compared four preprocessing strategies: (i) no preprocessing, (ii) StandardScaler, (iii) StandardScaler followed by PCA to 50 components, and (iv) StandardScaler followed by PCA to 100 components.

### 2.5 Molecular representations and kernel choices

When using GPs, we need to convert chemical SMILES string with numerical feature vectors. An efficient molecular representation can reduce the complexity of the problem by capturing only relevant information. Capturing all the relevant structure and chemical information, maintaining low dimensionality and providing chemical intuition are the challenges that any representation method has to deal with. By using three different molecular representations, we explore different aspects each with their unique tradeoffs. We use **ECFP Fingerprints**, i.e., Extended-Connectivity Fingerprints with radius 4, consisting of 4,096 binary features [33]. **MACCS Keys**, with 166-bit binary fingerprints representing predefined molecular fragments [34], and, **ChemBERTa Embeddings**, generated using the pre-trained ChemBERTa-77M-MTR model [35].

The choice of kernel function is fundamental to the GP’s ability to model correlations between data points based on their similarity. We explore five distinct covariance kernel functions viz., **Tanimoto, Linear, Radial Basis Function (RBF), Rational Quadratic (RQ)**, and **Matérn (***v* = 1.5).Please refer to the Supplementary Information (SI) for further details. For all kernels that include hyperparameters (i.e., Linear, RBF, RQ, and Matérn), these parameters (e.g., lengthscale *ℓ*, shape parameter *α*, outputscale *s*, and noise variance 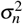) were optimized by maximizing the marginal log-likelihood during model training [36, 37]. For further information please refer to SI.

### 2.6 Model Explainability with SHAP

We incorporated SHapley Additive exPlanations (SHAP) [29, 38] to quantify the contribution of individual molecular features to GP model predictions across active learning cycles. For a molecule *x*, the prediction *f* (*x*) is decomposed into a baseline *φ*_0_ and additive contributions from *M* features,

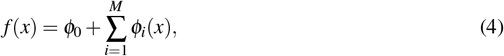

where *φ*_*i*_(*x*) denotes the SHAP value for feature *i*. Feature importance was computed as the mean absolute SHAP value across a held-out test set of *N*_test_ molecules,

Importance

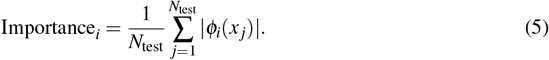

For each AL cycle, SHAP values were evaluated on 100 test molecules randomly sampled from the unqueried pool, using a background of 50 randomly sampled compounds from the training set to initialise the `shap.KernelExplainer`. The top ten features ranked by mean absolute SHAP value were retained for detailed analysis.The stability and robustness of these feature attributions were validated through quantitative analysis across different acquisition protocols.

For models trained on ECFP fingerprints, selected features were mapped back to molecular fragments using `RDKit`. **To address the ambiguity of mapping ECFP bits (due to bit collisions or multiple environments), we implemented an affinity-prioritized algorithm**. Atom environments corresponding to top-ranked fingerprint bits were first identified in all molecules containing the bit. These molecules were then sorted by descending affinity. The environment from the highest-affinity compound was extracted using `Chem.FindAtomEnvironmentOfRadiusN`, canonicalised to a SMILES string, and used as the representative fragment. These fragments were then ranked by a combined score of frequency and SHAP magnitude. This procedure ensures the identified chemical substructures are those most strongly associated with high-potency predictive signal and allows for a mechanistic interpretation of how AL reshapes the model’s representation of structure–activity relationships.

### 2.7 Experimental setup and evaluation

In order to evaluate the AL setup, we follow the fixed cost approach by Gorantla et al. [15] in acquiring a total of 360 compounds for each individual experiment. Each experiment starts with 60 randomly selected compounds, followed by 10 cycles of selecting 30 new compounds per cycle, using different exploration/exploitation strategies.

The cycle is then repeated for each experiment, and parameter combinations undergo repeated cycles. Suitable steps for updating and acquisition are undertaken to allow for unbiased comparison across dataset.

In this work, a single “experiment” refers to one complete, 10-cycle active learning simulation for a specific combination of dataset, molecular representation, kernel, acquisition protocol, and random seed.

For each dataset-representation-kernel combination, all seven acquisition strategies were evaluated, resulting in a total of 4 *×* 3 *×* 5 *×* 7 = 420 distinct experiments. The vast scope of the experiments poses a challenge to visualise and evaluate these results.

The entire computational study, including the training of all GP models, required approximately **4 hours** of wall-clock time on a single NVIDIA RTX 4090 GPU. This demonstrates the practical feasibility of applying our comprehensive benchmarking framework.

**Recall of Top Compounds (***R*_*k*_) metric quantifies the fraction of truly high-affinity compounds (top *k*%) that are successfully identified by the active learning process, relative to the total number of such compounds present in the entire dataset. It is calculated using the following equation 6,

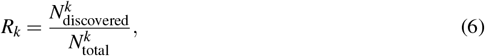

Where 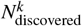 the number of compounds found in the acquired set that belong to the top *k*% class, and 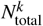 is the total number of compounds that actually belong in the top *k*% most active ones based on the observed activity in the entire dataset. Recall was computed for top 2% (*R*_2_) and 5% (*R*_5_) of compounds.

To provide a more comprehensive and robust assessment of early enrichment performance, we also report two additional standard metrics. The **Enrichment Factor (EF**_*k*_) measures how many times more frequently active compounds are found within the top *k*% of a ranked list compared to a random selection. It is defined as:

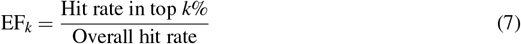

An EF_*k*_ of 1.0 corresponds to random performance. In this study, we report EF at 1%, 2%, and 5%.

To mitigate the sensitivity to a fixed cutoff *k*, we also report the **Boltzmann-Enhanced Discrimination of ROC (BEDROC)** score [39]. BEDROC is a metric that preferentially rewards the identification of active compounds at the top of a ranked list without requiring an arbitrary cutoff. It applies an exponential weight to each compound based on its rank, such that hits at the beginning of the list contribute much more to the final score than those found later. Following common practice for virtual screening, we use an *α* parameter of 20.0, which heavily focuses the evaluation on the top portion of the ranked list. The score ranges from 0 (no enrichment over random) to 1 (perfect ranking).

### 2.8 Data for the study

The active learning framework has been evaluated using four diverse protein target datasets viz, TYK2 (Tyrosine Kinase 2) [9], USP7 (Ubiquitin Specific Peptidase 7) [40], D2R (Dopamine D2 Receptor) [41], and MPRO (SARS-CoV-2 Main Protease) [26]. It is important to note that while Thompson et al. [9] describes TYK2 as a congeneric series derived from a single synthetic scaffold, our analysis using RDKit’s Murcko decomposition identified 104 distinct Murcko scaffolds, reflecting minor structural variations within the series. In contrast, USP7, D2R, and MPRO demonstrate substantially higher diversity (*N/M ≈* 0.41-0.45), reflecting more structurally varied compound collections. Details of datasets are provided in SI. Table 2 summarises some relevant details across the four datasets used.

**Table 2.** Dataset Properties.

| Property | TYK2 | USP7 | D2R | MPRO |
| --- | --- | --- | --- | --- |
| Binding Measure | pK<sub>i</sub> | pIC<sub>50</sub> | pK<sub>i</sub> | pIC<sub>50</sub> |
| Ligands (Total) | 9997 | 1799 | 2502 | 2062 |
| Scaffolds (Unique) | 104 | 770 | 1034 | 934 |
| Std Dev (p-value) | 1.36 | 1.31 | 1.44 | 0.91 |
| N/M ratio | 0.0104 | 0.428 | 0.413 | 0.452 |

We note that the datasets employ different affinity measures (pKi for TYK2 and D2R; pIC50 for USP7 and MPRO), as shown in Table 2. As these units are derived from different assay types and are not directly comparable, our study does not make direct, quantitative comparisons of the absolute affinity values across targets. Instead, our primary performance metric, Recall of Top Compounds (*R*_*k*_), is based on a relative, percentile-based threshold. For each dataset, the “top k%” active compounds are determined by internally ranking the molecules based on their specific affinity measure. This approach allows for a valid comparison of the *enrichment efficiency* of the AL strategies across the different chemical landscapes, without relying on a comparison of the raw activity scales.

## 3 Results and Discussion

While the conceptual idea of an active learning cycle is quite straightforward, the myriad of choices that one can make around surrogate models, acquisition functions, kernel choices, and molecular representations poses a challenge. Finding an optimal combination of choices may not be practical and evaluating the increasingly large number of combinations is difficult to assess and visualise. Lastly, active learning cycles are often black box systems allowing for little explainability of what the models are learning. Combining active learning with a SHapley Additive exPlanations (SHAP) [29, 38] analysis can provide some indications of model learning. With our results, we highlight that optimal AL strategies are highly context-dependent, underscoring the critical influence of inherent dataset characteristics and the complex interactions among methodological choices. In the following section, we present a comprehensive study of dataset characteristics followed by an analysis of how different methodological choices influence active learning performance. Furthermore, we explore how a versatile web-based tool aids in understanding complex results. Lastly, we use SHAP to understand if the AL cycles pick up patterns that lead to explainable properties that could be harnessed by medicinal chemist in designing effective AL strategies.

### 3.1 Chemical landscape sets difficulty scaffold diversity patterns anticipate AL headroom

We evaluated four therapeutically relevant targets with distinct chemistry—TYK2, USP7, D2R, and MPRO—to probe how dataset composition shapes active learning (AL) outcomes.

Scaffold diversity, as determined by the ratio of unique scaffolds to total molecules (*N/M*) is the main differentiator between the datasets. TYK2 exhibits exceptionally low diversity (*N/M ≈* 0.01), indicating a highly constrained chemical space dominated by few structural motifs. On the other hand, USP7, D2R, and MPRO exhibit significantly greater diversity (*N/M ≈* 0.41-0.45), which is indicative of more structurally diverse compound collections.

Scaffold diversity directly impacts molecular similarity patterns within each dataset as evident in figure 2. For instance, TYK2’s constrained chemical space is particularly evident with ECFP fingerprints, which show highly skewed similarity distributions with the majority of compound pairs exhibiting low Tanimoto similarities as evident from figure 2A. ChemBERTa embeddings and MACCS, on the other hand, display broader distributions centered at higher similarity values as evident from figure 2B, and C demonstrating how different representations highlight structural homogeneity differently. In contrast, USP7, D2R, and MPRO show wider and more diverse internal similarity distributions across all three molecular representations—ECFP, MACCS, and ChemBERTa (Figure 2 A, B, C). ECFP fingerprints produce sharp peaks at low similarity values, whereas MACCS keys and ChemBERTa embeddings give more spread-out distributions because they capture molecular structure in different ways.

**Figure 1.**
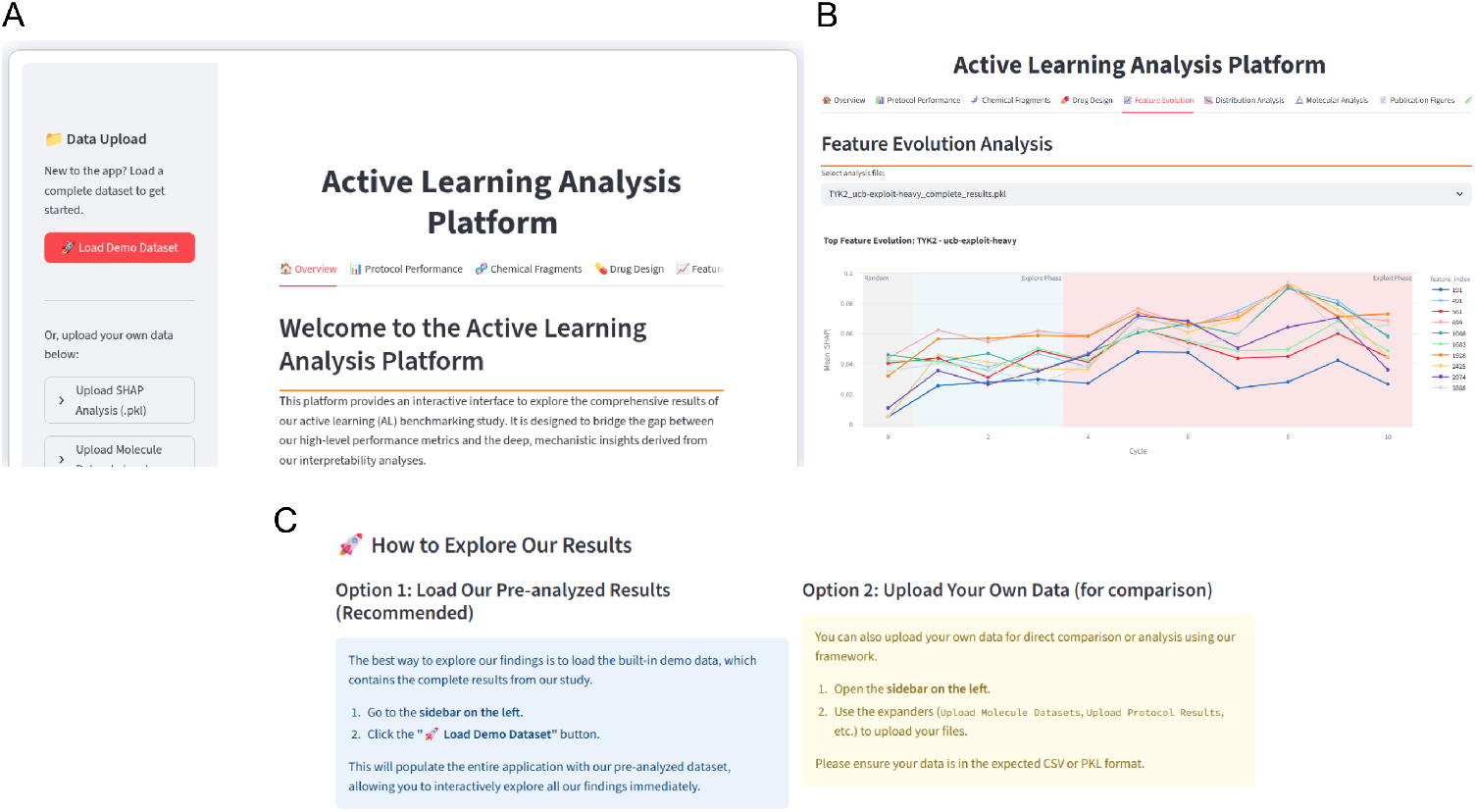
Screenshots of the interactive Active Learning Analysis Platform. (A) The main landing page, which offers two primary ways to engage with the tool: loading a complete, pre-analyzed demo dataset or uploading custom data files via the sidebar. (B) A view of the ‘Feature Evolution Analysis’ tab, which visualizes how the importance of top molecular features, as measured by their mean SHAP values, changes dynamically across the different phases (Random, Explore, Exploit) of the active learning cycles. (C) The “How to Explore Our Results” section, providing clear, step-by-step instructions for users to either explore the platform’s built-in findings or analyze their own data for comparison..

**Figure 2.**
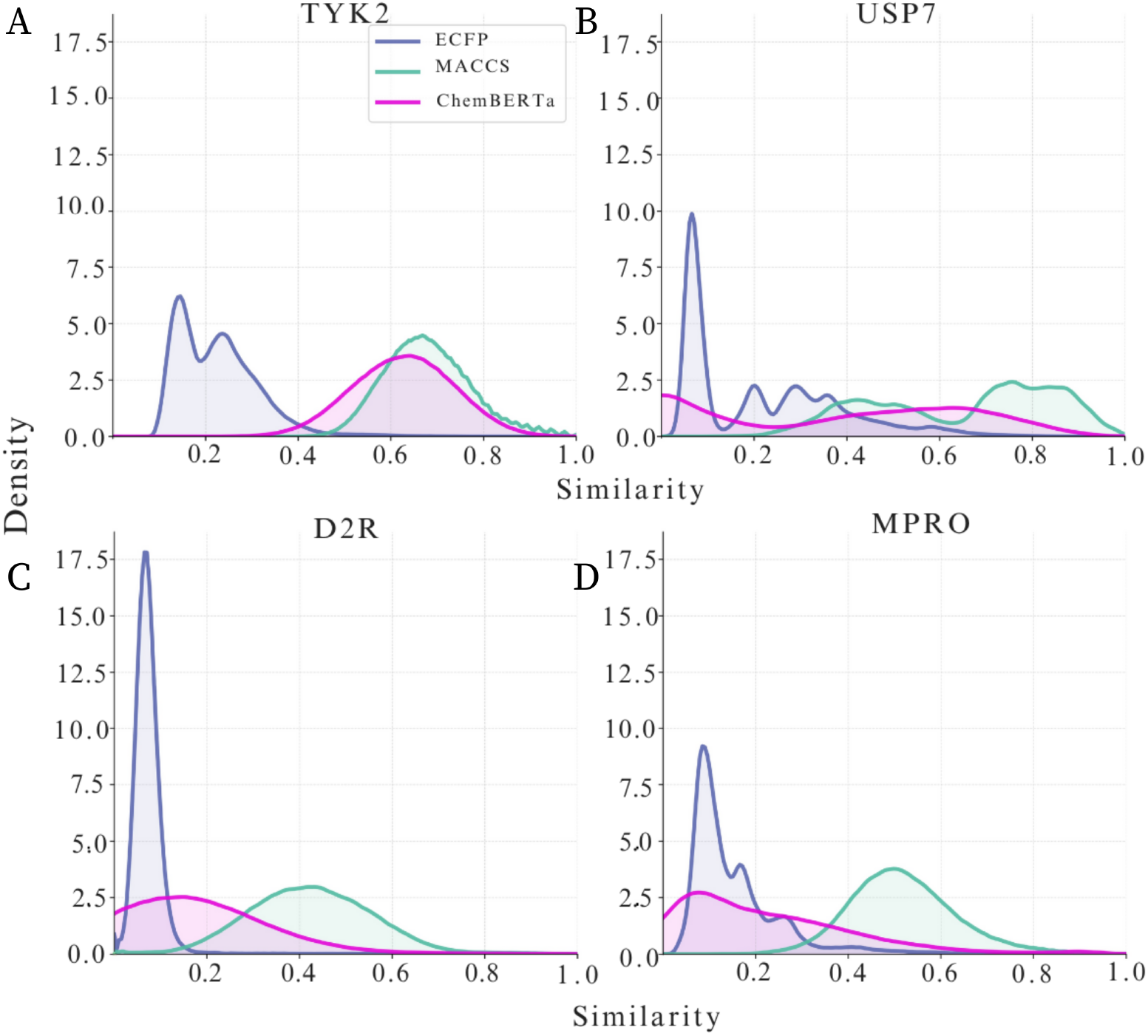
**Molecular Similarity Distributions Across Datasets and Representations. Kernel Density Estimate (KDE) plots illustrate the distribution of pairwise Tanimoto similarity scores for compounds within the TYK2, D2R, MPRO and USP7 datasets, as perceived by different molecular representations. For each dataset, the similarity profiles generated by ECFP4, MACCS, and ChemBERTa are compared. The ECFP fingerprints consistently show distributions heavily skewed towards low similarity across all datasets, particularly for TYK2, D2R, and MPRO. In contrast, both MACCS keys and ChemBERTa embeddings provide broader similarity distributions, often centered at higher values, indicating their capacity to capture more diverse structural relationships than ECFP.**

Dataset diversity patterns have direct implications for active learning performance. While the more expansive chemical landscape of USP7, D2R, and MPRO offers more chance of strategic compound selection, constrained chemical space like TYK2 restricts the opportunity for diversified exploration. Further dataset diagnostics are provided in the SI.

### 3.2 AL analysis platform

With 420 experiments run, the complexity of our results, which include four datasets, three molecular representations, five kernels, and seven protocols, necessitates a new way of presenting the results beyond graphs. To address this and promote transparency and reproducibility, we have developed the *Active Learning Analysis Platform*, an interactive web tool which is freely accessible. As shown in **Fig. 1**, this platform offers access to all the comprehensive experiments and analysis we did on each target dataset, enabling researchers to explore our findings interactively.

### 3.3 Dataset characteristics drive performance variation

The chemical space properties have a significant effect on active learning performance. With Recall of Top Compounds *R*_*k*_ values ranging from 0.5052 for the constrained TYK2 dataset to 0.9942 for the more diverse MPRO dataset indicating that performance varies significantly across datasets. Visual summary of the distribution of all experimental outcome with every dataset is available at https://shapanalysis.streamlit.app/.

Statistical analysis demonstrates that the intrinsic properties of the target dataset are the most dominant factor in determining achievable performance. To quantify the relative contributions of our methodological choices, we conducted a four-factor ANOVA (Type II Sums of Squares) on the final recall (Rk) values from all non-random protocols. The full model explained a substantial proportion of the variance in performance (*R*^2^=0.84,Adjusted *R*^2^ = 0.82).

To properly assess effect sizes, we computed omega-squared (*ω*^2^), an unbiased estimator of the population effect size, along with 95% bootstrap confidence intervals (1,000 iterations). Dataset identity exhibited the largest effect (*ω*^2^ = 0.31, 95% CI [0.28, 0.35]; F(3, 994) = 640.37, p < 0.001), confirming that the chemical landscape sets fundamental performance constraints. Notably, the interaction between dataset and kernel interaction showed a similarly large effect (*ω*^2^ = 0.31; F(12, 994) = 160.22, p < 0.001), demonstrating that kernel effectiveness is highly context-dependent.

Other factors made smaller but significant contributions: kernel choice (*ω*^2^ = 0.09, 95% CI [0.07, 0.11]; F(4, 994) = 135.27, p < 0.001), molecular representation (*ω*^2^ = 0.03, 95% CI [0.02, 0.04]; F(2, 994) = 91.47, p < 0.001), and the kernel × fingerprint interaction (*ω*^2^ = 0.04; F(8, 994) = 29.08, p < 0.001). The acquisition protocol, while statistically significant (F(5, 994) = 12.44, p < 0.001), had the smallest main effect (*ω*^2^ = 0.01, 95% CI [0.004, 0.019]), suggesting its role is to modulate outcomes within the constraints imposed by the dataset and model architecture. This statistical evidence reinforces that optimal active learning strategies are highly context-dependent, with dataset characteristics and their interactions with methodological choices playing the dominant role.

The Post-hoc Tukey HSD analysis showed that all UCB-based protocols performed significantly better than random selection in terms of mean Recall of Top Compounds *R*_*k*_ with all adjusted p-values less than 0.05, indicating strong statistical significance. However, there is no significant difference between the UCB protocols themselves, as all adjusted p-values were greater than 0.05. The practical impact of these improvements is measured using Cohen’s d effect sizes, which were larger, ranging from 0.934 `ucb-balanced` vs random to 1.308 `ucb-explore-heavy` vs random, revealing that UCB strategies had a strong advantage over random selection.

No one set of Kernel function, acquisition technique or molecular representation worked optimally in every circumstance. The best configuration for each dataset highlights the range of possible *R*_*k*_ values from 0.5052 for TYK2 to 0.9942 for MPRO, indicating that different datasets require different optimal setups.

### 3.4 Impact of Molecular Representation and Kernel Functions

Performance is significantly impacted by the kernel function and selected molecular representation. Our findings demonstrate no universally optimal combination, consistent with significant partial 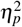 values for Dataset:Kernel interaction (65.92%) and Kernel:Fingerprint interaction (18.97%) in the ANOVA analysis.

#### Molecular Representations

ECFP fingerprints exhibited the most consistent and strong performance. Mean 2% Recall of Top Compounds (*R*_*k*_) of 0.37 *±* 0.31 across all datasets and protocols. ECFP demonstrated strong performance across a range of dataset-kernel combinations, particularly excelling in USP7 and MPRO with mean *R*_*k*_ values of 0.57 *±* 0.33 and 0.49 *±* 0.33, respectively. While ChemBERTa occasionally outperformed ECFP on specific combinations, ECFP provided superior predictability and delivered consistently reasonable performance even when other approaches yielded less than satisfactory results on challenging datasets like D2R and TYK2.

ChemBERTa embeddings exhibited a high-variance performance profile characterized by exceptional peaks and notable failures. When optimally paired with non-linear kernels i.e Matérn and RBF on USP7 and MPRO, ChemBERTa achieved the highest individual *R*_*k*_ of 0.99 on MPRO. This representation proved susceptible to significant performance loss under suboptimal conditions. On challenging datasets viz. D2R and TYK2, identical kernel combinations yielded dramatically lower mean *R*_*k*_ values, with some as low as 0.02 *±* 0.01 and a mean BEDROC of 0.003 ± 0.01 for the Matérn kernel on TYK2, highlighting ChemBERTa’s context-dependency and unpredictable efficacy.

MACCS fingerprints demonstrated the most consistent performance profile despite achieving the lowest overall mean *R*_*k*_ of 0.27 ± 0.18. This representation exhibited remarkably stable performance across different datasets, with substantially lower inter-dataset variance compared to ECFP or ChemBERTa. Even while MACCS rarely reached peak performance, its consistency makes it a reliable baseline when predictable results are prioritized over maximum performance. Notably, MACCS achieved competitive performance on D2R with *R*_*k*_ = 0.61 when paired with the Tanimoto kernel, demonstrating its potential for specific dataset-kernel synergies.

#### Kernel Functions

The Matérn and RBF kernels have been observed with the highest performance potential albeit a significant dataset-dependent variability. These kernels achieved the study’s peak *R*_*k*_ values of 0.9942 for MPRO with Matérn, and 0.97 for USP7 with Matérn when conditions were favorable, particularly with ChemBERTa or ECFP on receptive datasets viz. MPRO and USP7. For instance, MPRO with RBFKernel had a mean *R*_*k*_ of 0.75 ± 0.31, and USP7 with RBFKernel had 0.72 ± 0.33, a mean BEDROC of 0.6 ± 0.3, and a mean **Enrichment Factor at 2% (EF**_2_**) of** 27.9 ± 21.3. Conversely, these same kernels performed appallingly on challenging datasets, with TYK2 yielding mean *R*_*k*_ values as low as 0.04 ± 0.04, a mean BEDROC near zero (0.003 ± 0.01) and an EF_2_ of approximately 1.1 ± 0.8 on (Matérn) and 0.03 ± 0.02 (RBF), clearly showing their high-risk, high-reward characteristics.

The Linear and Tanimoto kernels delivered consistent, moderate performance across all tested conditions. Linear kernel achieved mean *R*_*k*_ of 0.35 ± 0.14 on D2R and 0.29 ± 0.13 on TYK2, and a mean **Enrichment Factor at 2% (EF**_2_**) of** 17.1 ± 8.2. This EF_2_ value, indicating that the top 2% of compounds were identified at over 17 times the rate of random selection, stands in stark contrast to the near-random performance of the non-linear kernels on the same dataset (EF_2_ *≈* 1.1), while Tanimoto kernel yielded 0.30 ± 0.12 and 0.26 ± 0.12 on the same datasets, respectively. These kernels maintained stable performance regardless of dataset difficulty or molecular representation.

The Rational Quadratic (RQ) kernel consistently underperformed across all conditions, achieving *R*_*k*_ as low as 0.12 ± 0.07, and EF_2_ of only 7.6 ± 4.0, on TYK2 and reaching only 0.26 ± 0.13 on MPRO. This demonstrates a trade-off wherein the non-linear kernels can offer high rewards but withhigh variability, while linear kernels offer reliable, moderate performance suitable for risk-averse applications.

#### 3.4.1 Impact of Active Learning Protocol

The active learning protocol had a considerable impact on both the trajectory and final outcome of the compound acquisition process, with distinct behavioural patterns observed across various dataset characteristics and kernel-representation combinations. While random selection consistently yielded the lowest performance (overall mean 2% Recall of Top Compounds (*R*_*k*_) of 0.11 ± 0.05 and mean EF_2_ of 6.4 ± 5.7), UCB-based strategies demonstrated clear advantages. Acquisition trajectories typically exhibited three distinct phases: an early exploration phase viz. 0–100 compounds, a middle transition phase with 100–250 compounds, and a late convergence phase with 250+ compounds.

Exploit-heavy strategies such as `UCB-exploit-heavy`, often designed for rapid prioritization, demonstrated effectiveness on USP7 and MPRO datasets, leading to rapid initial gains. Temporal SHAP analysis, which demonstrated top features for USP7 exploit-heavy strategies consistently peaking early in Cycles 2 or 3, indicates rapid initial SAR identification. In contrast,exploit-heavy strategies exhibited a noticeable ‘late spike’ in feature importance on dataset such as TYK2, suggesting that important SAR features are not immediately apparent rather are revealed after focused, persistent sampling in specific, high-reward regions of the chemical space. This ‘late spike’ reflects the model’s attempt to progressively prioritize subtle features within a highly constrained or challenging SAR landscape.

On the other hand, explore-heavy strategies such as `UCB-explore-heavy` typical showed slower initial progress but could achieve higher long-term *R*_*k*_ on complex datasets like D2R, showing more consistent improvement patterns. This reflects a broader sampling approach and a more distributed learning of features across the chemical space, as evident by less pronounced temporal shifts in SHAP feature importance. This approach is advantageous where targets have more diffused SAR or where novel active regions need to be discovered beyond narrow, pre-defined areas. Balanced and adaptive protocols (e.g.,`UCB-balanced, UCB-gradual`) frequently achieved competitive performance and demonstrated robustness across varied complexities, providing reliable options when optimal configurations are not immediately apparent.

The importance of protocol choice varied significantly depending on dataset selected. High-performing combinations such as `Matérn + ChemBERTa` achieved high *R*_*k*_ across most protocols with rapid convergence on datasets such as MPRO and USP7. On the other hand, protocol selection was more crucial for difficult dataset such as TYK2 and D2R which had significant *R*_*k*_ variation and demonstrated slow improvement beyond 300 compounds. This emphasizes how AL strategy effectiveness is highly dependent on dataset characteristics and the chosen kernel-fingerprint combinations, influencing initial trajectory and overall performance outcome.

### 3.5 Mechanistic Insights from Feature Importance Analysis

To obtain deeper mechanistic insights into how Gaussian Process models predict compound activity and how Active Learning influences the understanding of SAR we perform explainability studies. The application of SHAP analysis on ECFP fingerprint models is a well-established method [38] for understanding explainability. This analysis, focusing on TYK2 and USP7 targets uses exploit-heavy and explore-heavy AL protocols, to uncover distinct aspects of the model’s learning and the underlying chemical determinants of activity.

SHAP analysis consistently identified specific, chemically interpretable molecular fragments that were highly predictive of binding affinity, validating the model’s ability to learn genuine SARs [42, 43]. Importantly, compounds containing these top-ranked features consistently exhibited high binding affinities (Figure 5).

**Figure 3.**
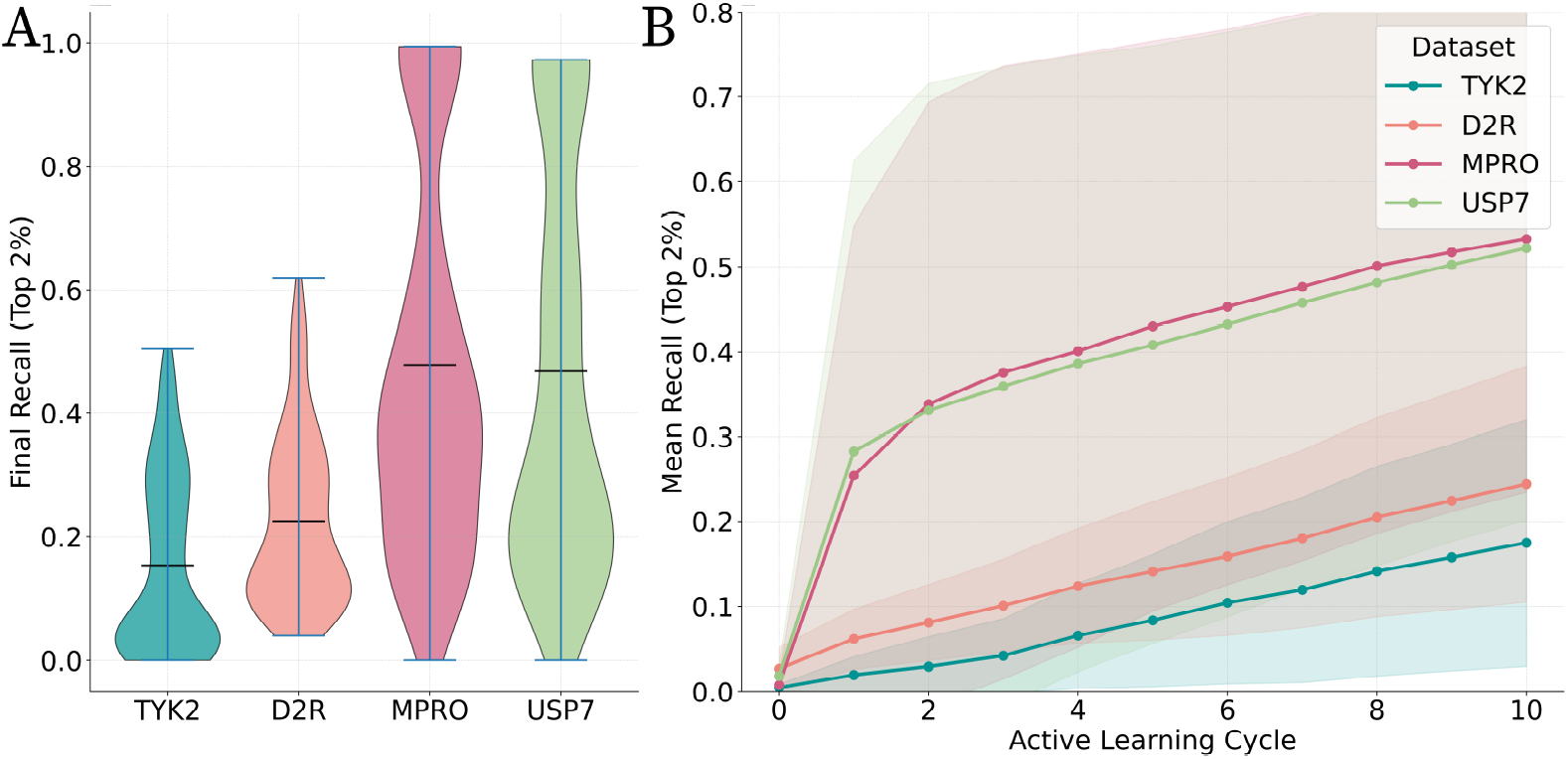
Overall Active Learning Performance Across Datasets. This composite figure summarizes key active learning performance metrics for each dataset, aggregating results across all kernel, molecular representation, and acquisition strategy combinations. **(A) Performance Distribution Across Datasets:** Violin plots illustrating the distribution of final 2% Recall of Top Compounds (*R*_*k*_) values for each dataset (TYK2,USP7, MPRO,D2R). The horizontal lines within each violin indicate the mean (*µ*) and median (red) *R*_*k*_ values, while the shape reflects the density of results. **(B) Learning Curves by Dataset:** Average 2% Recall of Top Compounds (*R*_*k*_) over the 10 active learning cycles, demonstrating performance evolution for each dataset. All plots aggregate data across all method combinations and replicates unless otherwise specified.

**Figure 4.**
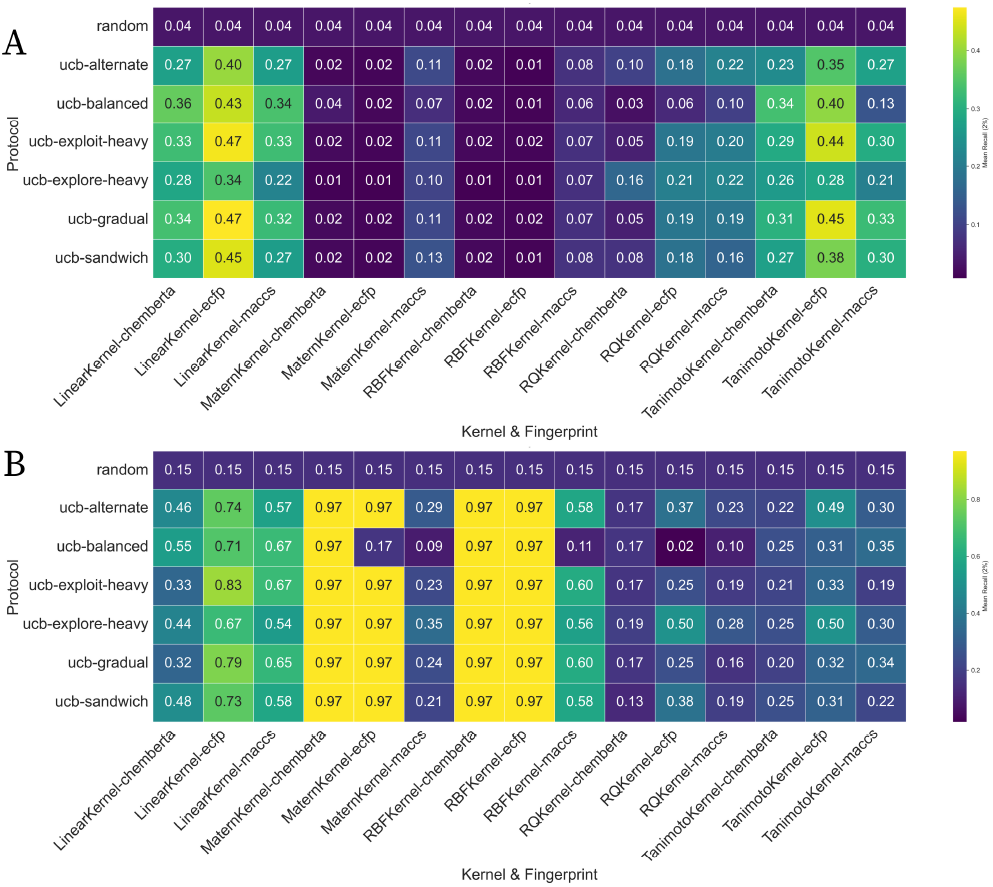
Mean 2% Recall of Top Compounds (*R*_*k*_) Across Protocols, Kernels, and Representations. Heatmap illustrating the average final *R*_*k*_ for each combination of active learning protocol (rows), Gaussian Process kernel (main columns), and molecular representation (sub-columns) at Cycle 10. Each cell represents the mean *R*_*k*_ across 3 replicate runs. The colour scale indicates performance, from low (dark purple/blue) to high (yellow). **(A) TYK2 Dataset:** Performance landscape for the challenging TYK2 dataset. Highlights the relatively lower overall *R*_*k*_ and the best-performing combinations. **(B) USP7 Dataset:** Performance landscape for the receptive USP7 dataset. Illustrates the generally higher *R*_*k*_ values and identifies highly effective combinations.

**Figure 5.**
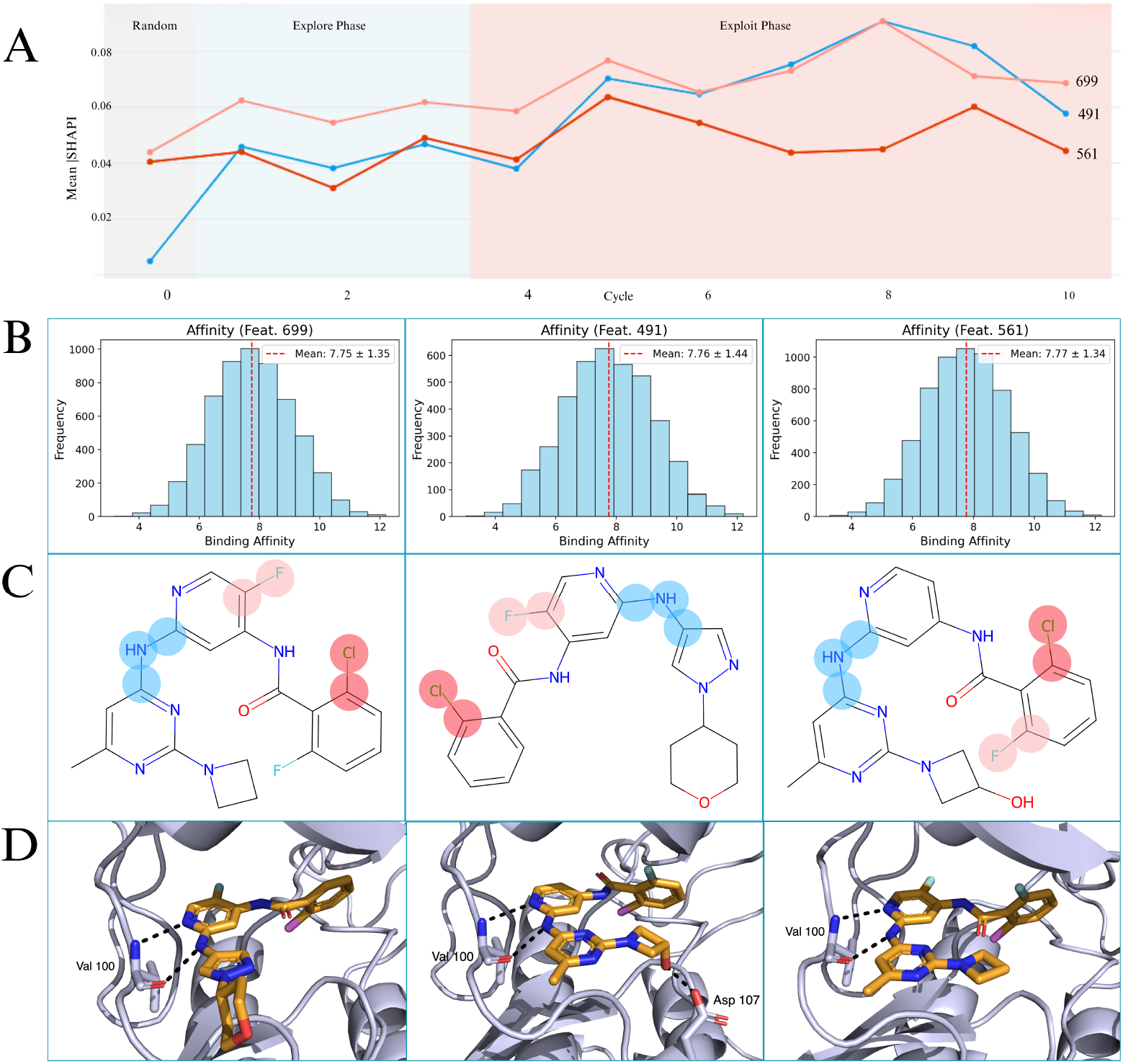
SHAP Analysis Reveals Dynamic Feature Importance in TYK2 Active Learning. **(A)** Mean SHAP importance evolution for top-ranking ECFP features (699, 491, 561) across active learning cycles using UCB-Exploit-Heavy protocol. Feature importance shifts from exploration (cycles 1–3) to exploitation phases (cycles 4–10). **(B)** Binding affinity distributions for compounds containing key features. Dashed lines show mean p*K*_*i*_ values: Feature 699 (7.75 ± 1.35), Feature 491 (7.76 ± 1.44), Feature 561 (7.77 ± 1.34), confirming association with high-affinity binders. **(C)** Representative TYK2 inhibitors with SHAP-identified substructures highlighted regions. Molecular structures demonstrate concrete chemical patterns underlying abstract feature importance scores. **(D)** Protein-ligand binding showing interaction modes for selected compounds in the TYK2 active site, with key residues Val100 and Asp107 labeled.

Our analysis demonstrates that the model learns stable and genuine SAR drivers. For the **USP7** target, the set of the top 5, most important features was identical between the `ucb-exploit-heavy` and `ucb-explore-heavy` protocols, yielding a Jaccard Index of 1.00. This perfect stability indicates that the model rapidly and consistently identified the core SAR. For the more challenging, low-diversity **TYK2** dataset, the analysis still showed good stability with a Jaccard Index of 0.43. While different protocols explored different nuances of the constrained chemical space, a core set of features (e.g., bits corresponding to cF and cNc fragments) were consistently ranked as most important. This provides strong evidence that our model is learning genuine SARs rather than stochastic noise.

#### Key Predictive Fragments and Chemical Relevance

For TYK2, key features consistently identified included across both exploit heavy and explore heavy methods included **halogenated motifs** — such as Feature ID 699, cF; Feature ID 561, cCl — and **nitrogen-containing aromatic systems** such as Feature ID 491, cNc; Feature ID 2425, ccc(nc)Nc. These features were repeatedly highlighted as significant determinants for TYK2 activity. These fragments with mean affinity of TYK2 6.76-7.78 p*K*_i_ align with common interaction modes for kinase inhibitors, such as halogen bonding and *π*-stacking [3, 44]. According to the chemical pattern analysis, TYK2’s primary characteristics included 100% aromatic, 54.8% halogen-containing, and 30.1% nitrogen-containing fragments.

For USP7, prominent features were consistently associated with **carbonyl groups** such as Feature ID 2362, C=O and **nitrogen-rich heterocycles** such as Feature ID 3500, cnc for both protocols.

These features with mean affinity for USP7 9.33-9.66 pIC50 are chemically relevant for deubiquitinase active sites, often involved in hydrogen bonding and electrostatic interactions [45, 46]. Further suggested by the identification of a complex fragment ID 875, i.e, `nc1cncn(CC2(O)CCNCC2)c1=O` suggests the model’s capability to prioritize intricate patterns. USP7’s top fragments were 100% aromatic, 24% nitrogen-containing, and 0% halogen-containing, aligning with DUB modulator characteristics.

#### Robustness of Insights Across Active Learning Protocols

The identified key features and their associated mean affinities remained remarkably consistent between exploit-heavy and explore-heavy AL protocols for both TYK2 and USP7. For instance, in TYK2, Feature ID 699 (cF) consistently ranked highest across both protocols, with identical affinity statistics. Similarly, for USP7, Feature ID 3500 (cnc) and Feature ID 2362 (C=O) maintained high ranks and consistent affinities across protocols. This robustness suggests that the identification of core binding motifs is stable, even if the sampling strategy influences the diversity of compounds explored around them [44]. This consistency provides further confidence in the model’s generalizability and its robust mechanistic understanding of binding, even when the underlying sampling strategies might aim for different balances of exploration and exploitation within the chemical space.

## 4 Conclusion and Outlook

In this work, we evaluated active learning (AL) strategies for ligand binding affinity prediction, investigating the interplay between molecular representations, kernel functions, and acquisition protocols across various chemical datasets. Our main conclusion is that AL’s effectiveness varies significantly based on the dataset’s chemical properties. Statistical analysis demonstrated that the dataset, and its interaction with techniques like kernel functions, is the primary factor influencing performance, establishing the limits for AL success.

Our analysis revealed important trade-offs between different methodological choices. We discovered that simpler, explicit representations like ECFP fingerprints, paired with robust linear kernels, offer consistent and reliable performance across a wide range of dataset complexities. On the other hand, advance, pre-trained embeddings like ChemBERTa, when combined with flexible non-linear kernels such as Matérna and RBF, can achieve state-of-the-art peak performance; however, they are prone to catastrophic failures on difficult or mismatched chemical landscapes. Similar to this it was demonstrated the the AL protocol selection is context-dependent. Exploit-heavy methods are better suited for rapid lead optimization within well-defined SARs, whereas explore-heavy strategies are beneficial for novel chemotype discovery in more diverse chemical spaces. Mechanistic insights from our SHAP analysis offers a framework for understanding why these choices matter, linking them to the model’s dynamic learning of SARs throughout the AL cycles.

According to these results, there is no “one-size-fits-all” AL strategy that works in all circumstances. We proposed a context-aware framework for AL in drug discovery demonstrating promising results in terms of ease of their analysis. Practitioners should first analyze their dataset’s chemical space, i.e., scaffold diversity and similarity to set reasonable expectations and select AL components accordingly. Challenging or unknown spaces may benefit from stable combinations such as ECFP with a Linear kernel, while well-behaved SARs might justify using risky, high-reward methods like ChemBERTa with non-linear kernels.

While this study provides a robust framework, it has limitations, including its retrospective nature and the focus of SHAP analysis on ECFP models. Future work can focus on the prospective validation of these findings in real-world drug discovery campaigns. The most promising future direction, however, lies in the development of adaptive Active Learning frameworks. These systems could learn the characteristics of the chemical space in real-time and automatically select or adjust the molecular representation, kernel, and acquisition strategy during the campaign, moving beyond the static protocol choices. We can fully utilize active learning to speed up the development of novel medicine by balancing the performance improvement in ligand binding affinity prediction with explainability built in the model from the start. Further improvements could also be achieved by exploring more advanced surrogate models, such as Warped Gaussian Processes, which could allow the model to explicitly learn the non-Gaussian distribution of affinity data.

## Acknowledgements

This work was supported by the Department of Computer Science and Engineering,Shiv Nadar University, Delhi-NCR, and the United Kingdom Research and Innovation (grant EP/S02431X/1), UKRI Centre for Doctoral Training in Biomedical AI at the University of Edinburgh, School of Informatics, and Exscientia Plc, Oxford.

